# Fully Automated Point Spread Function Analysis Using Pycalibrate

**DOI:** 10.1101/2022.10.11.511725

**Authors:** J Metz, M Gintoli, AD Corbett

## Abstract

Reproducibility is severely limited if instrument performance is assumed rather than measured. Within optical microscopy, instrument performance is typically measured using sub-resolution fluorescent beads. However, the process is performed infrequently as it is requires time and suitably trained staff to acquire and then process the bead images. Analysis software still requires the manual entry of imaging parameters. Human error from repeatedly typing these parameters can significantly affect the outcome of the analysis, rendering the results less reproducibile. To avoid this issue, PyCalibrate has been developed to fully automate the analysis of bead images. PyCalibrate can be accessed either by executing the Python code locally or via a user-friendly web portal to further improve accessibility when moving between locations and machines. PyCalibrate interfaces with the BioFormats library to make it compatible with a wide range of proprietary image formats. In this study, PyCalibrate analysis performance is directly compared with alternative free-access analysis software PSFj, MetroloJ QC and DayBook3 and is demonstrated to have equivalent performance but without the need for user supervision.

## 1. Introduction

As the diversity and complexity of microscopy techniques continues to increase, there is a greater focus on reproducibility and quality control (QC) to ensure claims made using the image data are justified (Hammer et al., 2021; Nelson et al., 2021). One of the key barriers to the regular QC of optical microscopes is the time required to conduct the highly skilled task of quantifying instrument performance. Automating the QC process is expected to both significantly reduces the time and the level of training required to complete the task.

Microscope QC has three main stages: (i) sample preparation, (ii) image acquisition and (iii) analysis. Variability is introduced at every stage. Using a standardised protocol, it can take several days to develop the bead sample (Cole et al., 2011). Once the sample has been produced it will have a limited30 lifetime (approx. 6-12 months). Fortunately, commercial, pre-prepared slides are now available e.g. PSF check slides (University of Exeter Consulting, Exeter, UK), Gatta-Beads(Gattaquant, Munich, Germany) or Tetraspeck slides (ThermoFisher Scientific, MA, USA) to accelerate the process and improve standardisation. Automation of image acquisition remains a challenge due to the proprietary nature of hardware control in commercial microscopes. This paper focuses on the automation of the analysis stage which, when conducted across multiple beads, image channels, objectives and microscopes, still presents a significant investment in time. Acquiring image stacks of sub-resolution beads gives direct access to the three-dimensional shape of the microscope point spread function (PSF) (Schneider & Webb, 1981), from which it is possible to extract parameters including spatial resolution, colour channel alignment and (for multiple beads in the field of view) field flatness. The axial and lateral full-width at half-maximum (FWHM) of the PSF can be compared with theoretical values to indicate the presence of chromatic aberrations, poor index-matching or inadequate objective lens cleaning. Measuring the PSF across the field of view can also reveal the presence of field-dependent aberrations.

There are several software packages available that obtain PSF FWHM from bead image data. Some are Java-based, provided as plugins to the ImageJ/Fiji platform (Schindelin et al., 2012) e.g. PSFj (Theer et al., 2014) and MetroloJ QC (Faklaris et al., 2022). Commercial choices such as Huygens (Scientific Volume Imaging, Hilversum, Netherlands) exist alongside semi-commercial options such as DayBook3 (Argolight, Bordeaux, France) which is free to use but incurs fees for data storage. Although popular, these products all heavily depend on the user to introduce or tweak parameters to perform an accurate analysis. This not only greatly increases the time required to perform the analysis but unconscious errors in these values will lead to potentially significant and unrecognised errors in the analysis results.

In this paper, we introduce fully automated software analysis, which not only reduces the time required for data analysis but significantly reduces the variability in the results of the analysis by eliminating human error. In the event that PyCalibrate is unable to retrieve the required information from the metadata, PyCalibrate highlights these values as missing and the resultant FWHM values will be provided in pixel values. As with the alternative software solutions, PyCalibrate can interpret a wide range of image formats (including czi, oir, lif, nd2), can analyse multiple beads (up to 200) across the field of view and provides analysis reports both in PDF and CSV format for direct integration into the lab workflow. Unlike the alternative solutions, PyCalibrate is also freely available via a web app, which stores the analysis reports, together with the raw data files on the cloud so that they are available to the user from anywhere in the world. PyCalibrate is also available as Python code which benefits from the convenience of license-free development tools, and the vast collection of libraries available for code extension and modification.

The following Results section provides an introduction to processing PSF data via the PyCalibrate web app. This is followed by a direct performance comparison with popular software tools (PSFj, MetroloJ QC and DayBook 3) when analysing identical synthetic and real data sets.

## 2. Results

### 2.1 Running PyCalibrate web app

PyCalibrate can be accessed through the web app at https://www.psfcheck.com/psfcheck-processing which also contains a tutorial and instructional video which are summarised below. PyCalibrate can also be accessed by downloading the Python code directly from https://gitlab.com/psfcheck/pycalibrate-psf.

#### 2.1.1 Using the PyCalibrate Web App

##### (i). Register an account

Navigate to https://www.psfcheck.com/ and click the “PyCalibrate” tab (Figure 1(A(i))). From the “Menu” option below, select “Register” (Figure 1(A(ii))). Enter your details and click “Register”. An email from will automatically be sent to the contact email address to enable the user to confirm their PyCalibrate account.

**Figure 1:**
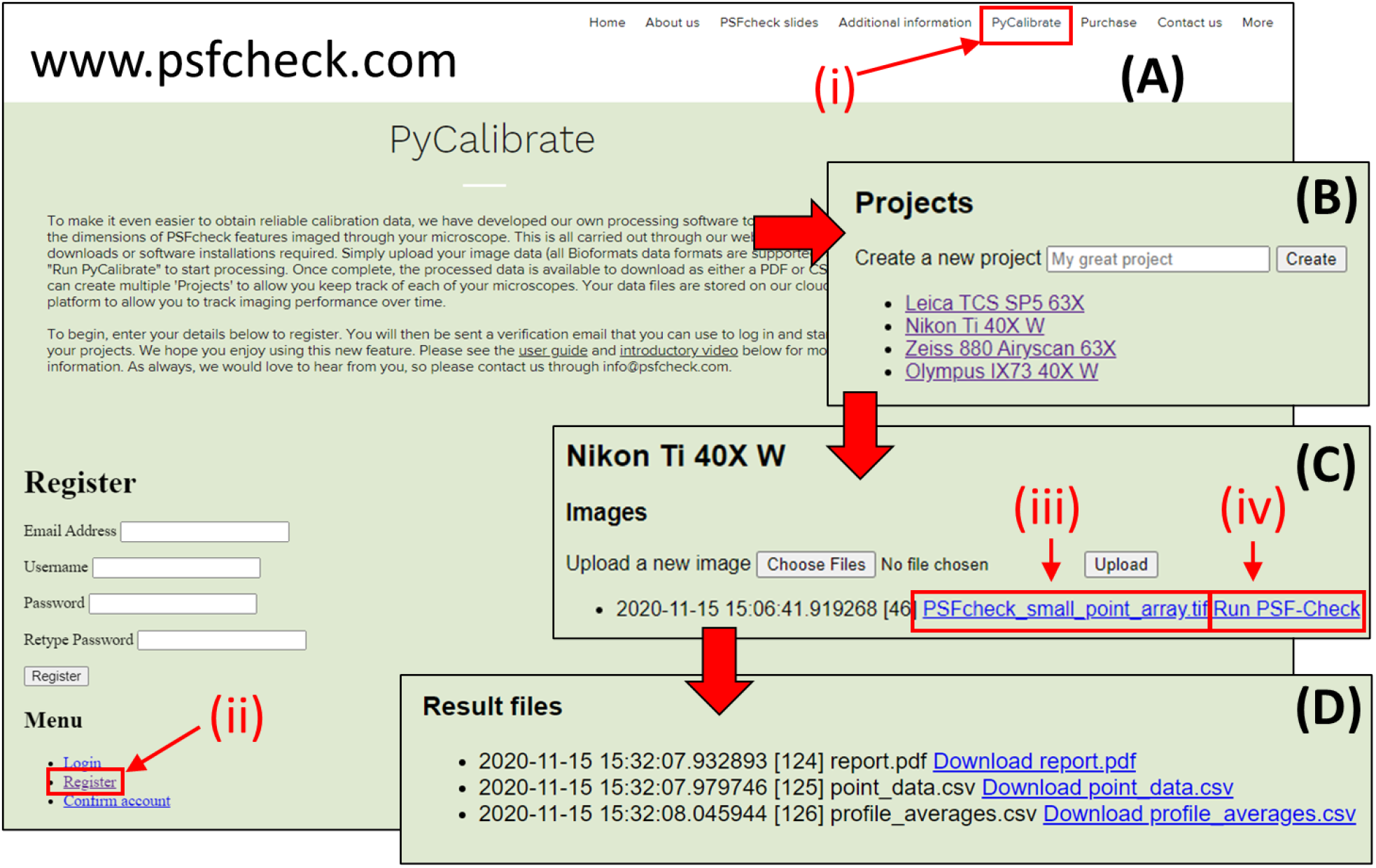
Four steps required to process data using PyCalibrate. After navigating to the website (A(i)) and registering an account (A(ii)), the user can create projects (B) in which to collect image stacks of point spread functions. Once an image stack has been uploaded (C) the user can either download the raw data (C(iii)) or process it (C(iv)). After processing, the PSF reports can be downloaded as an overview PDF or CSV files for either the average across all points, or all features individually (D).

##### (ii). Create a project

To help the user to organise calibration files, individual projects can be created. A project could be an individual microscope, or a specific microscope and objective lens combination (Figure 1(B))

##### (iii). Upload and process image data

Once a project has been created, click on “Choose files” to bring up a dialog box and select the image stack to be analysed. Once selected, clicking “Upload” to upload it to the cloud server. Once the file has uploaded, two links appear; the first has the name of the file just uploaded and the second is labelled “Run PSF-Check”. The raw data file can be downloaded again at any time by clicking the link with filename (Figure 1(C(iii))). This can be useful for reviewing raw data at a later date. Clicking “Run PSF-Check” (Figure 1(C(iv))) will run the PyCalibrate software on the raw data file.

Please note:

- The raw data file should only contain a single image stack (z-stack).
- If the data file contains multiple colour channels, a report will be generated only for the first colour channel (i.e. the lowest index in the colour channel stack).

##### (iv). Download results

After a few minutes of processing (depending upon file size), refresh the page to see a new “Results” link. If the results link has not appeared, the file has not yet completed processing. PyCalibrate offers three output files (Figure 1(D)). Each of these files can be downloaded directly by clicking on the link. The first is a PDF file summarising the appearance of the calibration data, average values across the field of view and the individual fitting values for each detected point. The second and third files are in CSV format, containing either fitting values for each of the individual points in the field of view or averages, according to the information the user requires.

The PDF file begins with filename of the raw data file together with a time and date stamp (Figure 2(C)). Also shown is an image of the raw data, showing a maximum intensity projection together with overlays showing which features have been detected and the order in which they appear in the “Raw fit data” table (Figure 2(B)).

**Figure 2:**
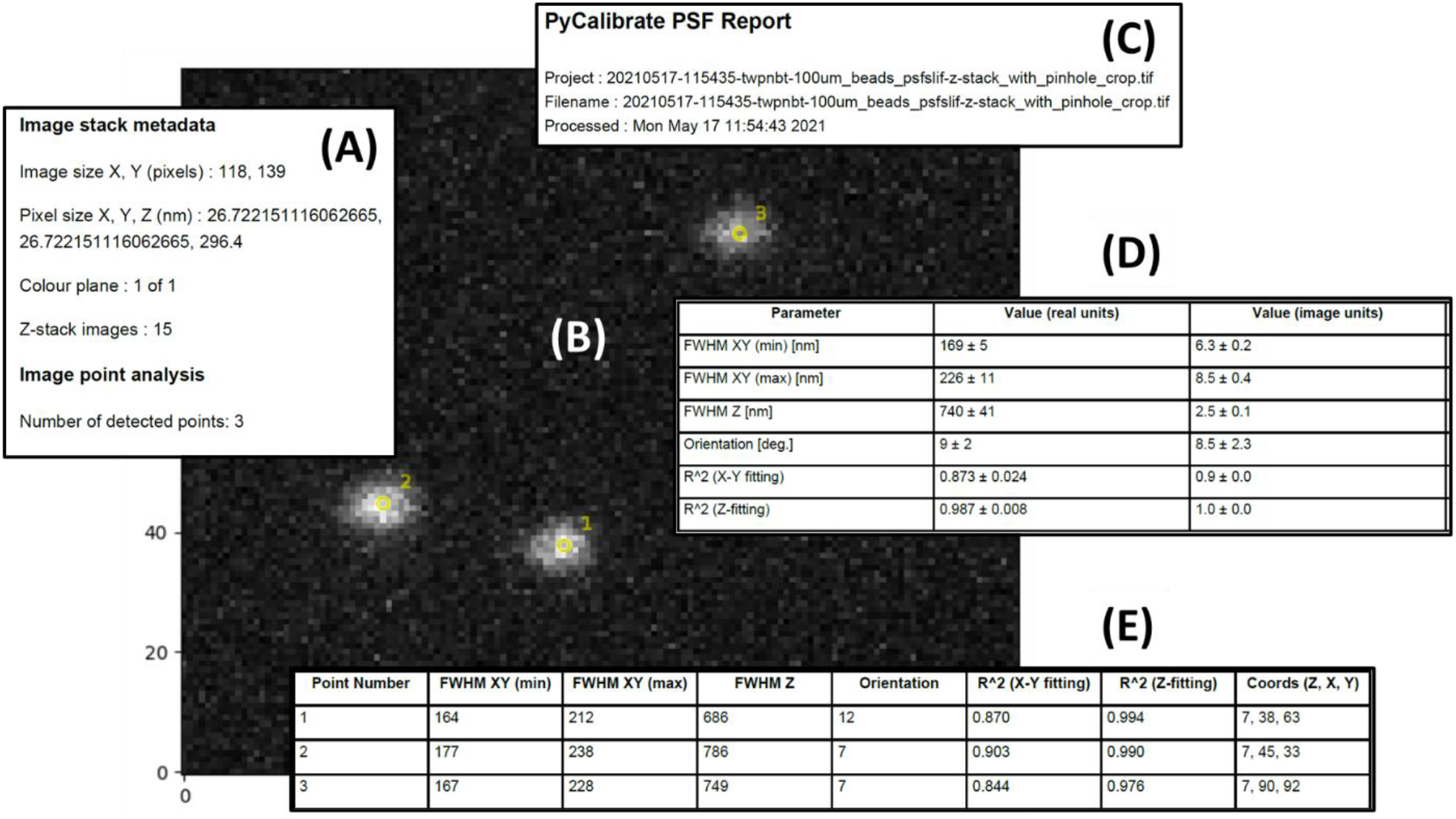
Key features of the PyCalibrate analysis results PDF summary. The PDF includes a description of the retrieved file metadata (A), a maximum intensity projection of the data set (B), a time stamp for the analysis together with the name of the file used (C), the lateral and axial dimensions of the detected features, both averaged across the field of view (D) and enumerated for each feature individually (E).

Please note:

- If PyCalibrate has not been able to successfully extract the X, Y or Z pixel dimensions from the image metadata, the parameter value will show as “NA”.

The PDF file continues with an “Average values” summary table containing the averages for all of the features detected across the field of view (Figure 2(D)). The first column shows the parameter measured, the second the average value in nm and the third column shows the average parameter value in pixels, together with ±1 (sigma) uncertainty values. The confidence in the fitting is given by the R^2^ values. These same parameters are provided for each individual point identified across the field of view in the “Raw fit data” table (Figure 2(E)).

Heat maps (not shown) are provided for each of the main fitting parameters. For densely populated fields (i.e. many detected points), this allows the user to quickly identify variations across the field of view.

### 2.2 Synthetic PSF data tests

Synthetic PSF data sets were generated, spanning a range of signal to background ratio (SBR) and effective pixels size (see Materials and Methods). The data sets were processed by PyCalibrate and each of the comparison software packages. In all cases efforts were made to get the best out of each comparison software package by manually adjusting threshold levels and regions of interest to encourage an accurate result. It is worth noting that whilst PSFj recommends an SBR ≥ 50 and DayBook 3 recommends an SBR ≥ 10, it is still possible to obtain measures of the spot widths at lower SBR values.

Unlike experimental data, the lateral and axial values of the synthetic FWHM are known absolutely. It is therefore possible to calculate the absolute error in the value returned by the analysis software. The lateral RMS error is calculated as the root mean square of the x-error and y-error, expressed as a percentage of the true width:

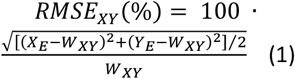

Where *X*_*E*_ and *Y*_*E*_ are the estimated FWHM along the x-axis and y-axis respectively by the analysis software and *W*_*XY*_ is the true width of 273 nm. Similarly, the root mean square error was calculated for the axial FWHM using:

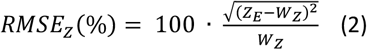

The lateral and axial RMSE(%) values are shown for each of the 32 synthetic data sets and for each of the software packages in Figure 3(A-D) and Figure 3(E-H) respectively. These tables use a heat map with five bands to indicate whether the RMSE(%)is <10%, 10-20%, 20-30%, 30-40% or >40%. In cases where the RMSE(%) is greater than 100%, the values were capped at 100%. In the event that no value was retuned from the software (analysis failed) the default RMSE(%) was set to 100%.

**Figure 3:**
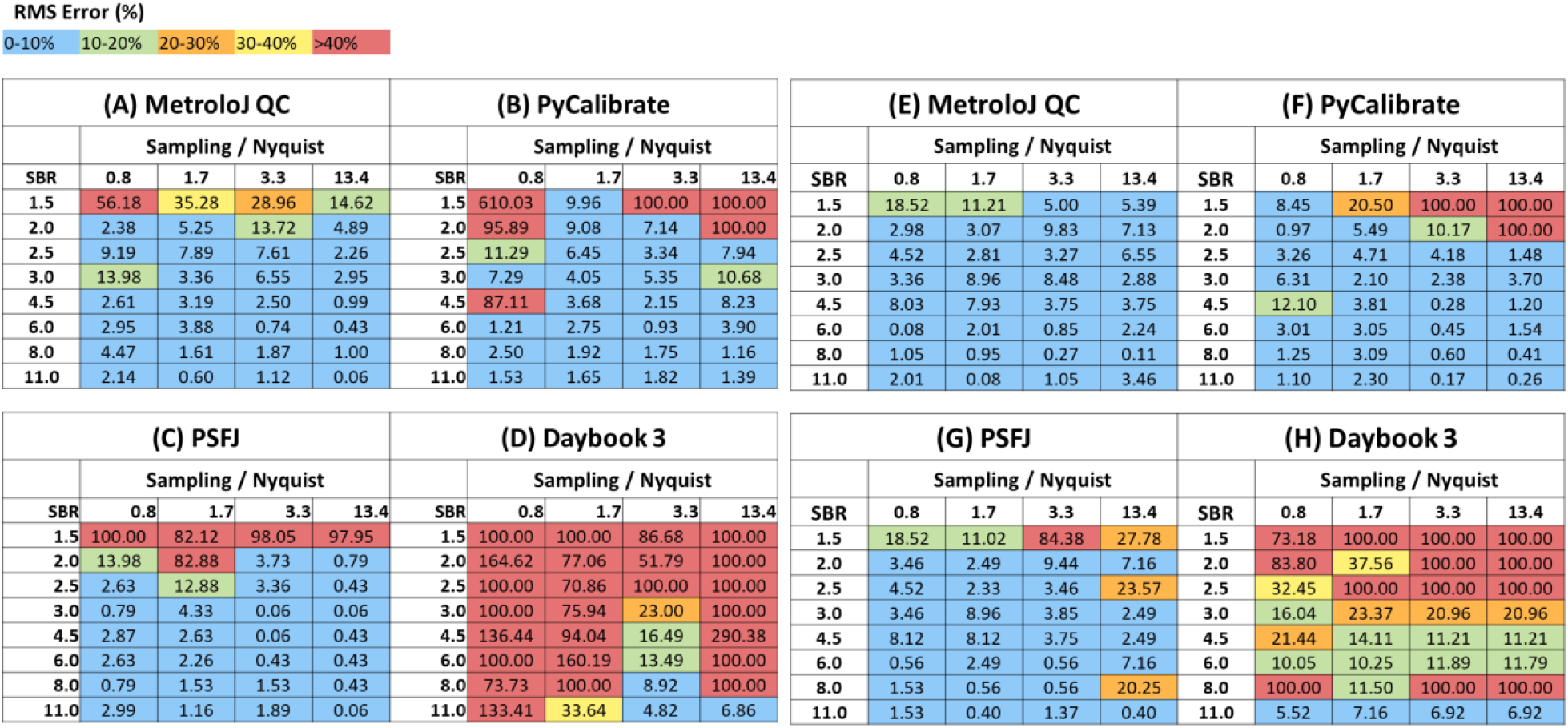
Comparison of the error in the determination of the lateral FWHM for synthetic data with variable sampling and signal to background ratio. The absolute root mean square error (RMSE) is expressed as a percentage of the true lateral FWHM (273 nm).

Overall, the lateral measurements indicate that the performance is similar for PyCalibrate, PSFj and MetroloJ QC (Figure 3(A-C)), with MetroloJ QC performing slightly better at the more challenging low SBR and coarse sampling data sets (towards the top left of the array). By contrast the Daybook 3 software struggled to perform as well as the other three packages. All software packages performed better when measuring the axial FWHM (Figure 3(E-H)). This is largely because changes in lateral sampling (effective pixel size) did not have a significant impact on the axial profile, as the axial sampling rate remained fixed for all data sets. Variation in performance can still be seen with SBR.

### 2.3 Experimental PSF data tests

Unlike the theoretical PSF data sets which were symmetric in X and Y, the experimental data sets may exhibit a degree of ellipticity. In this case it is important to acknowledge the differences between the values reported by the different software packages. PyCalibrate and PSFj provide maximum and minimum values for the lateral FWHM, corresponding to a 2D Gaussian fit where the perimeter delineating the extent of the spot is an ellipse. PyCalibrate and PSFj provide the semi-major and semi-minor axis measurements as well as the angle of the semi-major axis relative to the horizontal (X) image axis. MetroloJ QC and DayBook 3 provide the spot widths when projected along the X and Y dimensions of the image. To provide the most accurate comparison, it was necessary to map from the elliptical descriptions of the spot perimeter to the X and Y image axes. The mathematics to perform this mapping is described in the Materials and Methods section. This precaution was taken even though there was no clearly visible ellipticity present in the confocal images that would induce large degrees of ellipticity and the result of the mapping procedure was expected to be very subtle.

Image stacks were acquired of fluorescent features of increasing size and SBR (see Materials and Methods). The extracted X and Y widths of these features are shown in Figure 4(A) with the feature size projected along the X image axis shown as green bars and along the Y image axis as orange bars. This shows that for the first three packages (PyCalibrate, PSFj and MetroloJ QC) the results are broadly similar. However, as with the simulated data sets, DayBook 3 often provides highly inaccurate values, in many cases, the values returned being much larger than the vertical scale shown in Figure 4(A).

**Figure 4:**
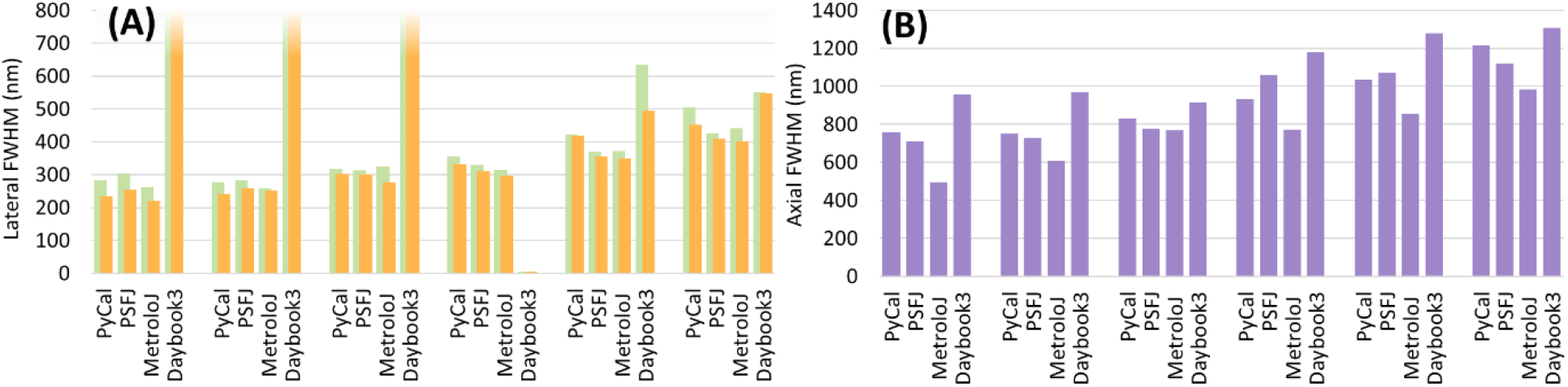
(A) Lateral (XY) sizes for each of the six PSFcheck feature sizes imaged. The measured feature widths along the x- and y-axes are shown in green and orange respectively for each of the analysis packages. The fluorescent feature size increases from left to right. (B) Axial FWHM for each of the six PSFcheck features measured by the four software packages.

For the axial FWHM (Figure 4(B)) the values are once again more comparable between the four packages. Box and whisker plots (Figure 5) are also provided to better illustrate the variability between the first three packages when applied to the same experimental image data. As there are only three values for each box, the upper, central and lower lines of the box correspond to the values from the three packages. The average value is shown as a black cross.

**Figure 5:**
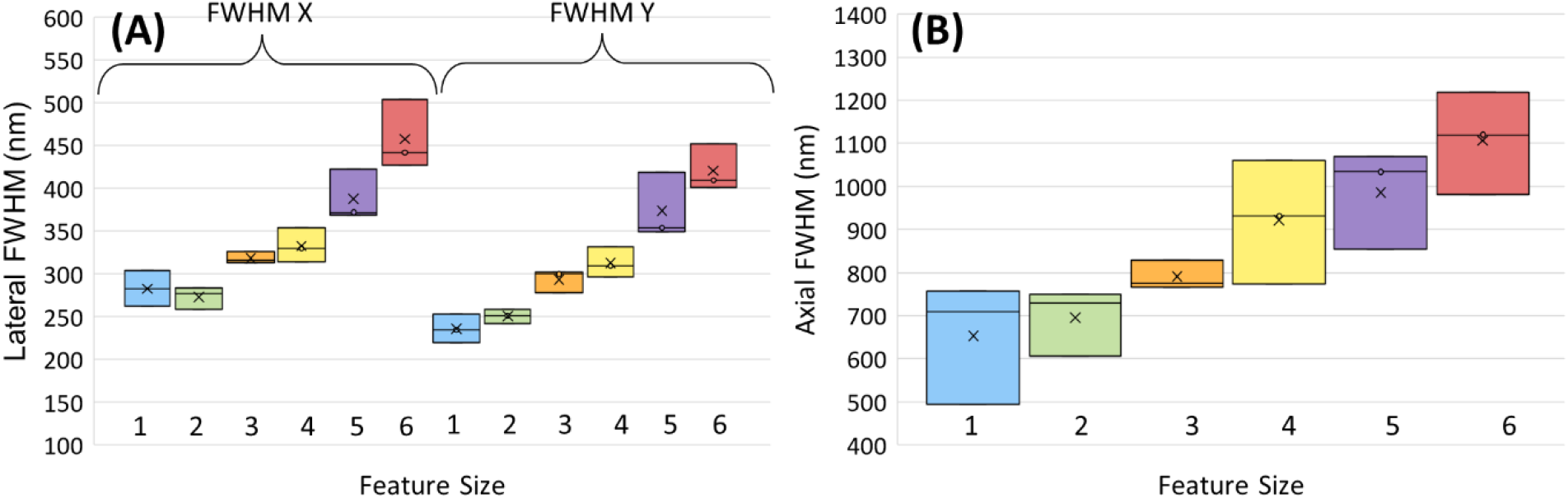
Box and whisker plots demonstrating the distribution of (A) lateral and (B) axial FWHM values determined by three of the four analysis software packages (PyCalibrate, PSFj and MetroloJ QC). The software packages analysed confocal image stacks of six different feature sizes in a PSFcheck slide.

Figure 5 demonstrates that the differences in the analysis of the identical data sets can be as high as 50 nm laterally and 200 nm axially. However, the standard deviation of the three values, averaged across all data sets, was just 17 nm laterally and 77 nm axially.

## 3. Discussion

PyCalibrate, a fully automated software tool for the analysis of PSF image data has been shown to produce results comparable to, or exceeding, leading alternative freeware solutions. Uniquely, PyCalibrate is available both as executable code and as a web app to further improve access and usability. The software doesn’t require manual user input, extracting values automatically from the image file metadata. The results of a comparison study using both synthetic and experimental data demonstrated that there were no systematic differences between the results obtained by PyCalibrate and those of PSFj and MetroloJ QC. The standard deviation of the results from the three packages had an average of 17 nm laterally and 77 nm axially. This corresponds to less than one pixel laterally and one quarter of a pixel axially. DayBook 3 struggled to provide accurate values for the given data sets, but this may be addressed in later revisions. Whilst there exist other differences between the software packages in terms of functionality, usability and speed, these were not examined here.

With full automation and greater accessibility through the web app, PyCalibrate will enable both new and experienced microscope users to obtain an accurate measurement of one of the most important parameters required for microscopy calibration.

## 4. Materials and Methods

### 4.1 Accessing comparison software

Instructions for accessing and operating the comparison software are available as follows. The PSFj software (build 231) is available with a manual from the “Supplementary Software” section of the corresponding publication: https://doi.org/10.1038/nmeth.3102.

MetroloJ QC .jar file (version 1.1.3) was accessed via https://github.com/MontpellierRessourcesImagerie/MetroloJ_QC. Finally, DayBook3 (version 1.8.5) was downloaded from https://argolight.com/document-and-share-results-with-daybook-software/. PDF documentation can be accessed via the software. There is also a video tutorial provided here: https://argolight.com/blog/how-to-use-the-point-spread-function-psf-analysis-in-daybook-webinar/.

### 4.2. Generating synthetic PSF data

All real FWHM measurements made on images of sub-resolution beads are estimates of the true PSF dimensions. This uncertainty makes it difficult to state exactly what error is introduced by the analysis software. To overcome this, synthetic data was created using a 3D Gaussian spot model. This is not an entirely accurate description as diffraction limited PSFs are in fact Airy functions and aberrated PSFs can adopt a wide variety of shapes. This shape was chosen because all the software packages map two- or three-dimensional Gaussian functions to the image data. By using a model 3D Gaussian to begin with we can then more closely assess their accuracy as a function of signal to background ratio (SBR) and effective pixel size, rather than their ability to map onto the many and varied non-Gaussian profiles observed in real microscopes.

The model 3D Gaussian was chosen to have lateral and axial FHWM of 273 nm and 1036 nm respectively. Whilst the choice of FWHM values is arbitrary, these values approximately correspond to the theoretical PSF of a 1.15 NA water immersion lens at an emission wavelength of 515 nm.

The FWHM values were converted into sigma values describing the Gaussian width using a conversion factor of 2ln(2) (≈2.36, (Theer et al., 2014)). Using the *σ*_*X,Y*_ and *σ*_*Z*_ value together with the amplitude (‘*A’*) of the Gaussian peak, intensity values were calculated for each voxel. A background offset (‘*B*’) was added to each of the values within the image volume. The value of B=100 counts was used for all synthetic data sets. The model assumed that the images are Poisson noise dominated. The intensity value of each voxel in the noise-free model was taken to be both the mean and variance values of a Poisson distribution. Voxel values in the noise-free model were then replaced by values selected at random from the corresponding Poisson distribution.

The model lateral FWHM (XY) is 273 nm. The maximum pixel size to achieve Nyquist sampling is therefore half of this value, 136.5 nm. The synthetic data sets used effective pixel sizes of 10 nm, 40 nm, 80 nm and 160 nm (T*able 1*). These values corresponded to sampling rates which varied from 0.8 × Nyquist to 13.4 × Nyquist. The axial sampling rate was kept the same for all data sets.

**Table 1:** The different levels of spatial sampling used for each model PSF

| Voxel size XY (nm) | XY Sampling (/Nyquist) | Voxel size Z (nm) | Z Sampling (/Nyquist) | 3D image size (pixels) |
| --- | --- | --- | --- | --- |
| 10 | 13.4 | 100 | 5.2 | 512 x 512 x 25 |
| 40 | 3.3 | 100 | 5.2 | 128 x 128 x 25 |
| 80 | 1.7 | 100 | 5.2 | 64 x 64 x 25 |
| 160 | 0.8 | 100 | 5.2 | 32 x 32 x 25 |

The amplitude values for the peak Gaussian value (‘A’) varied from 50 counts to 1000 counts (*Table 2*). The SBR was defined as *SBR* = (*A* + *B*)/*B* (*Table 2*), corresponding to SBR values ranging from 1.5 to 11. With four sampling rates for each of the eight SBR values produced 32 synthetic data sets overall, some of which are represented in Figure 6.

**Table 2:** Peak signal, background and SBR values used in each of the Gaussian models.

| Gaussian peak value, A (counts) | Background noise level, B (counts) | Signal to background ratio (SBR) |
| --- | --- | --- |
| 50 | 100 | 1.5 |
| 100 | 100 | 2 |
| 150 | 100 | 2.5 |
| 200 | 100 | 3 |
| 350 | 100 | 4.5 |
| 500 | 100 | 6 |
| 700 | 100 | 8 |
| 1000 | 100 | 11 |

**Figure 6:**
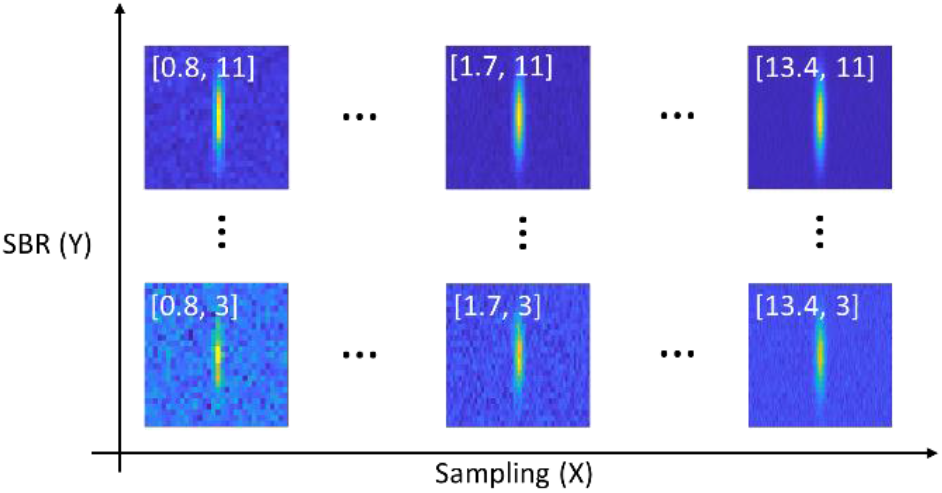
XZ slices through six of the 32 synthetic data sets used to test the analysis software. The X coordinate shows the effect of increasing the lateral (XY) sampling rate from 0.8 × Nyquist to 13.4 × Nyquist. The axial sampling rate was kept the same for all data sets. The Y coordinate shows the impact of increasing the signal to background ratio (SBR) from 3 to 11.

### 4.3 Experimental PSF data

To assess performance using real data, fluorescent features of variable size (and hence SBR) were imaged on a Leica SP5 confocal microscope using a 63X 1.4 NA oil objective (details below). The fluorescent features were created by direct laser writing on a PSFcheck slide (Corbett et al., 2018). The PSFcheck slide contains five different patterns. In this work, the ‘single point power series’ pattern was used (Figure 8). This pattern consists of rows of fluorescent features, with feature size changing between rows. The rows alternate between features written with single femtosecond pulses (smaller) and five femtosecond pulses (larger). The feature size of the single pulse features decreases monotonically with the row number (Figure 8). These features were chosen as they allowed a more continuous variation in feature size (and SBR) than bead sizes available commercially.

All data sets were acquired on a Leica TCS 5 using an oil index 63X 1.4 NA objective. Samples were illuminated at 488 nm (< 0.1 mW) with fluorescent emission collected over a broad spectral window (500-750 nm). 1024 × 1024 × 15 voxel images were collected at 200 Hz sampling rate using a four-line average. All data sets used the same voxel size (24 nm × 24 nm × 300 nm voxels). The image of an individual spot was extracted by cropping in XY to a 3 μm × 3 μm region of interest. Software packages analysed a single image stack at each feature size to compare performance as a function of SNR.

### 4.4 Calculating the x- and y-projections of an ellipse

To ensure a fair comparison between software packages, it is necessary to map from min/max FWHM values provided by PyCalibrate and PSFJ to the X/Y FWHM values provided by MetroloJ QC and Daybook3, (there is insufficient information for the reverse mapping). PyCalibrate and PSFJ provide max/min and theta values, corresponding to the semi-major and semi-minor axes of an ellipse inclined at an angle of theta to the x-axis. Projecting the limits of an inclined ellipse onto the x- and y-axes requires a formula which is derived below.

Figure 7 shows a sketch of the ellipse inclined at an angle theta from the x-axis. The values of *a* and *b* represent half of the calculated maximum and minimum FWHM values respectively. It can be seen in this case the errors introduced by calculating the projections using *a* · cos *θ* and *b* · sin *θ* for the x-axis and y-axis projections. For a more accurate calculation, we seek to first determine *X*^′^ and *Y*^′^ (shown in blue). These can be easily calculated if the angles *ε*_*x*_ and *ε*_*y*_ are known, together with the chord lengths *k*_*x*_ and *k*_*y*_ using the following formulae:

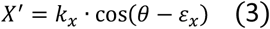

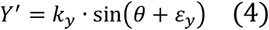

**Figure 7:**
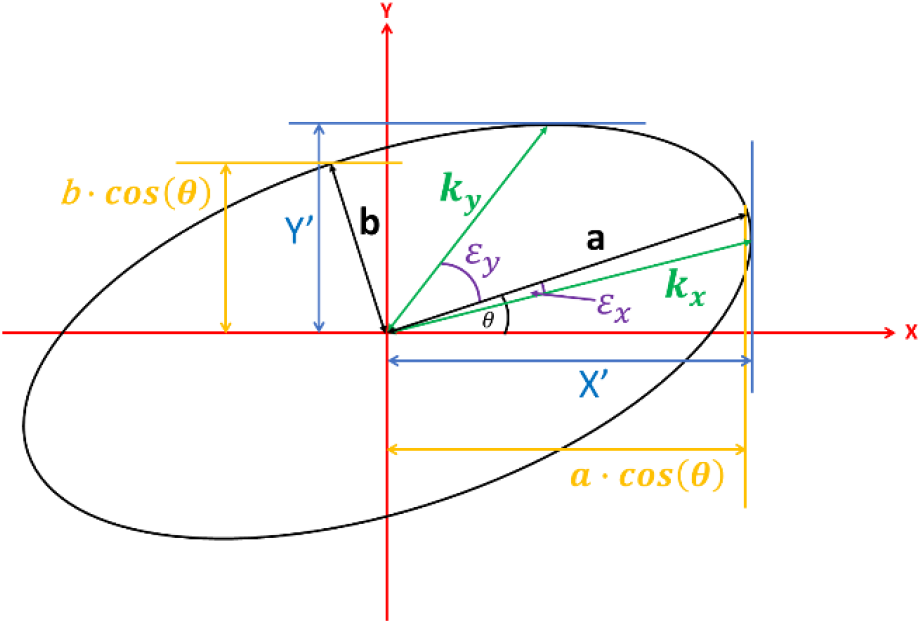
Parameters used to calculate the proper lateral and vertical extent of a rotated ellipse.

The chord lengths, *k*_*x*_ and *k*_*y*_ can be determined from *ε*_*x*_ and *ε*_*y*_ using the standard formula for the chord length of an ellipse:

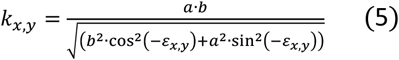

To calculate the angles *ε*_*x*_ and *ε*_*y*_, we consider the rotation of the ellipse by an angle θ in the clockwise direction (Figure 9).

**Figure 8:**
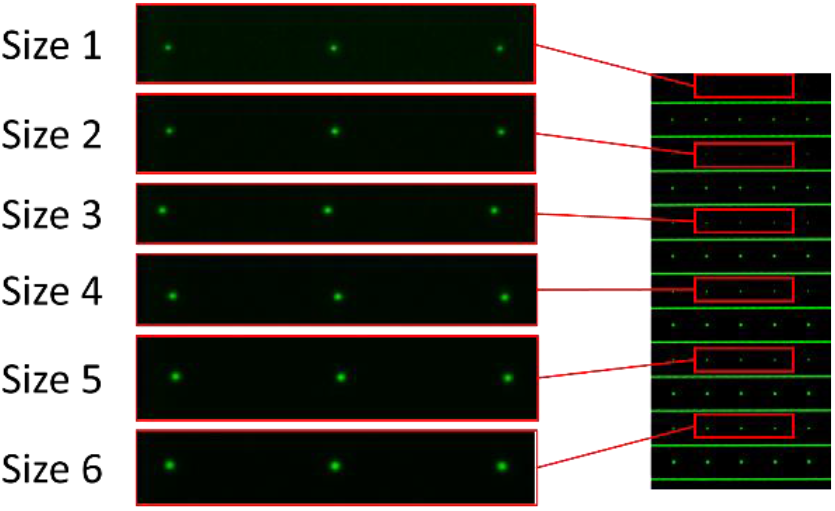
Fluorescence images of PSFcheck slide features (left) taken from alternate rows of the single point power series pattern (right).

**Figure 9:**
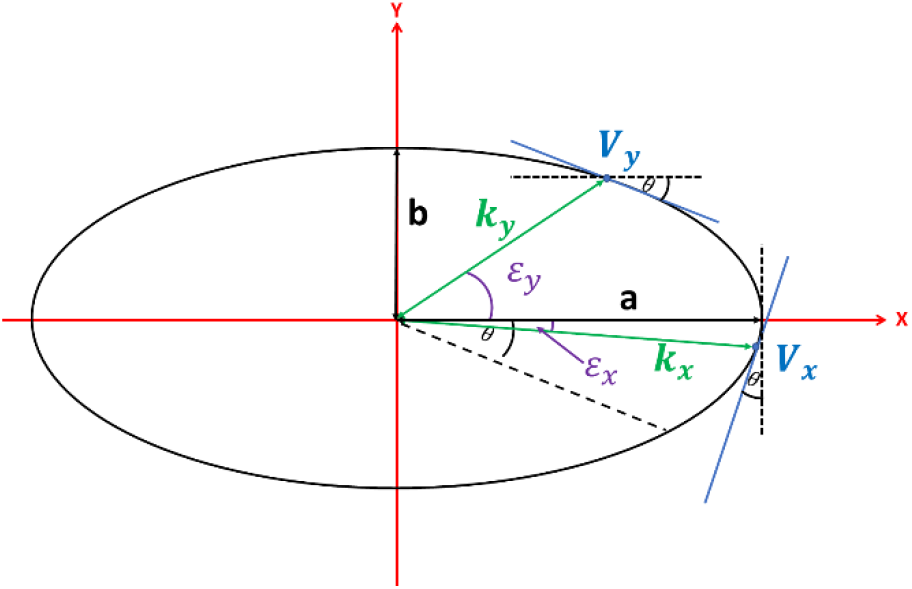
Clockwise rotation of the ellipse in Figure 7 by angle θ

The gradients (*m*_*x*_ and *m*_*y*_) at each vertex (*V*_*x*_ and *V*_*y*_) on the rotated ellipse (Figure 9) are known to be *cot*(*θ*) at *V*_*x*_ and −*tan*(*θ*) at *V*_*y*_. Comparing these known gradient values to the standard expressions for the gradient of an ellipse, we can uniquely identify *ε*_*x*_ and *ε*_*y*_:

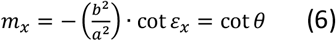

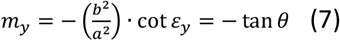

As *a, b* and *θ* are known, we can calculate *ε*_*x*_ and *ε*_*y*_ from equations (4) and (5). Using equation (3) we can calculate *k*_*x*_ and *k*_*y*_. Finally, using equations (1) and (2) we can calculate the projections *X*^′^ and *Y*^′^.

We now have all of the information required to perform the mapping for *θ* in the range [0, *π*/2]. We can see by symmetry that a negative value of *θ* in the range [0, −*π*/2]will produce the same result for the projection. This allows the modulus to be taken for *θ* values in the range [−*π*/2, *π*/2]. For |*θ*| > *π*/2, we require a mapping that monotonically decreases *θ* again until we reach zero at the value of |*θ*| = *π*. This can be achieved using the following mapping:

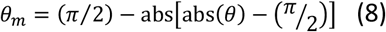

## Data Availability

The research data supporting this publication are openly available from the University of Exeter’s institutional repository at: https://doi.org/10.24378/exe.XXXX (TBC).

